# A vibrating mesh nebulizer as an alternative to the Collison 3-jet nebulizer for infectious disease aerobiology

**DOI:** 10.1101/594358

**Authors:** Jennifer D. Bowling, Katherine J. O’ Malley, William B. Klimstra, Amy L. Hartman, Douglas S. Reed

**Affiliations:** Center for Vaccine Research, University of Pittsburgh, Pennsylvania, USA; Department of Immunology, University of Pittsburgh, Pennsylvania, USA; Department of Infectious Diseases and Microbiology, University of Pittsburgh, Pennsylvania, USA

**Author notes:** Address correspondence to Douglas S. Reed.

## Abstract

Experimental infection of animals via inhalation containing pathogenic agents is essential to understanding the natural history and pathogenesis of infectious disease as well as evaluation of potential medical countermeasures. We evaluated whether the Aeroneb, a vibrating mesh nebulizer, would serve as an alternative to the Collison, the ‘gold standard’ for generating infectious bioaerosols. While the Collison possesses desirable properties that have contributed to its longevity in infectious disease aerobiology, concerns have lingered about the volume and concentration of agent required to cause disease and the damage that jet nebulization causes to the agent. For viruses, the ratio of aerosol concentration to nebulizer concentration (spray factor, SF), the Aeroneb was superior to the Collison for four different viruses in a nonhuman primate head-only exposure chamber. Aerosol concentration of influenza was higher relative to fluorescein for the Aeroneb compared to the Collison, suggesting that the Aeroneb was less harsh to viral pathogens than the Collison when generating aerosols. The Aeroneb did not improve the aerosol SF for a vegetative bacterium, *Francisella tularensis.* Environmental parameters collected during the aerosols indicated that the Aeroneb generated a higher relative humidity in exposure chambers while not affecting other environmental parameters. Aerosol mass median aerodynamic diameter was generally larger and more disperse for aerosols generated by the Aeroneb than what is seen with the Collison but ≥80% were within the range that would reach the lower respiratory tract and alveolar regions. These data suggest that for viral pathogens, the Aeroneb is a suitable alternative to the Collison 3-jet nebulizer.

**Importance:** The threat of aerosolization is often not the natural method of transmission. While selection of an appropriate animal model is vital for these types of experiments, other confounding factors can be controlled through a thorough understanding of experimental design and the effects that different parameters can have on disease outcome. Route of administration, particle size, and dose are all factors which can affect disease progression and need to be controlled. Aerosol research methods and equipment need to be well characterized to optimize the development of animal models for bioterrorism agents.

## Introduction

Experimental infection of animals with aerosolized pathogens to study pathogenesis or evaluate medical countermeasures remains a complicated procedure that requires expert training and highly sophisticated equipment. Environmental and situational factors can affect the survival, dose, site of deposition, and virulence of pathogenic agents (1–4). For example, studies have shown that relative humidity inside the chamber can alter aerosolization of bacteria and viruses (3, 5–7). Particle size can affect where a pathogen lands in the respiratory tract, which can have dramatic effects on pathogenesis and virulence (1, 2, 4, 8, 9). Therefore, to achieve reproducible dosing between experiments, one must fully characterize and validate all parameters of an aerosol exposure.

The Collison 3-jet nebulizer is a commonly employed aerosol generator in infectious disease aerobiology research (Fig. 1A). The nebulizer utilizes Bernoulli’s principle to shear a liquid suspension into aerosolized particles, which impact against a hard surface (the interior of the jar) to further break apart particles (10). A primary reason for the appeal of the Collison nebulizer is that it generates high concentrations of particles that are relatively monodisperse with a mass median aerodynamic diameter between 1-2 μm (11). This particle size can reach the alveolar regions of the lung. However, some studies suggest the shear forces, impaction, and recirculation of the infectious sample can damage organisms, potentially reducing pathogen viability or infectivity (12, 13). Damaged bacteria or viruses may also stimulate immune responses that protect the host. These effects could raise the dose required to cause disease, thereby requiring large quantities of pathogens grown to high titers for aerosol experiments. While the process of aerosolization will always place mechanical stress on infectious agents, aerosol generators that are ‘gentler’ than the Collison would be desirable.

**Figure 1.**
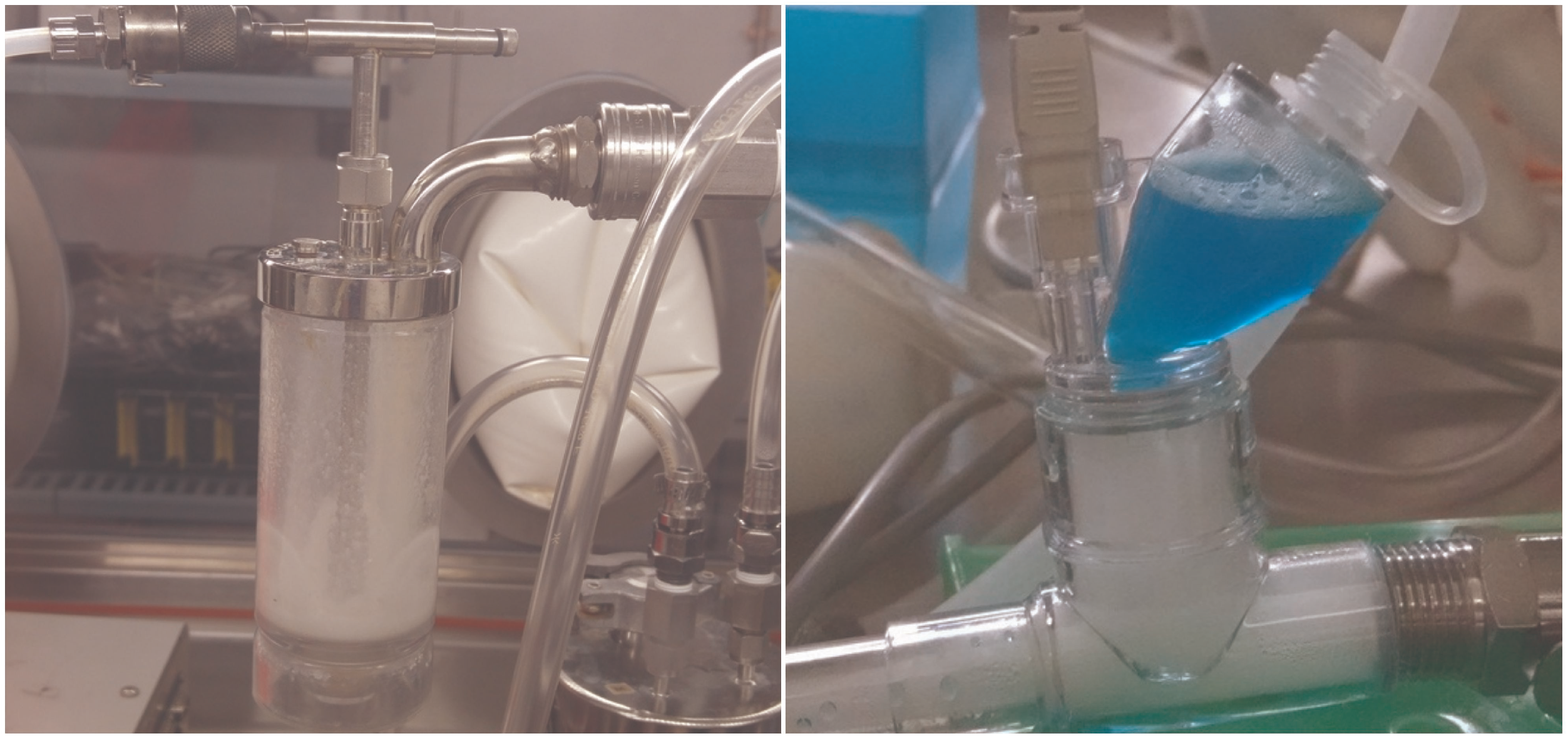
Collison and Aeroneb aerosol generators. The Collison nebulizer by CH technology (left) utilizes Bernoulli’s principle to create aerosols from a recirculated liquid sample. The Aeroneb by Aerogen (right) utilizes a vibrating palladium mesh membrane to create aerosols from a liquid sample.

The Aerogen Solo (a.k.a Aeroneb) is a single-use nebulizer employed in clinical settings for the delivery of aerosolized medication. The Aeroneb utilizes a palladium mesh perforated with conical shaped holes that act as a micropump when vibrated rather than high velocity air flow (14). We hypothesized that the Aeroneb might be gentler on pathogens than the Collison, potentially leading to improved aerosol performance. In this report, we report our efforts to characterize the aerosol performance of the Aeroneb as compared to the Collison for representative bacterial *(Francisella tularensis)* and viral pathogens (influenza).

## Results

### Aerosolization of viruses

Experimental aerosolization of pathogenic agents is commonly evaluated by determination of the spray factor (SF), which is calculated as the ratio of the aerosol concentration to the starting concentration. This allows one to compare between different aerosols to evaluate the impact of aerosol generators, sampling devices, and environmental parameters. A less commonly used alternative is aerosol efficiency (AE) that compares the amount of agent aerosolized to what is recovered from aerosol sampling devices. Prior to comparing nebulizers, we first sought to determine whether there was a difference in aerosol performance of H3N2 and H1N1 influenza viruses in the ferret whole-body (FWB) and rodent whole-body (RWB) chambers with the Collison nebulizer (Figure 2). No significant differences were seen between the SF of H1N1 and H3N2, regardless of the chamber used. Influenza spray factors (SFs) were slightly higher in the RWB compared to the FWB but this difference was also not statistically significant. The H1N1 data included aerosols with A/Ca/4/09 or A/PR/8/34; no significant differences existed between the two isolates based on a two-sided Mann-Whitney test (*p* =0.0901). In other experiments using other chambers and nebulizers, no differences in SF were seen based on the choice of influenza subtype, strain, or method of propagation (eggs or cell culture); the results in Table 1 and Figure 3 show combined results for all influenza viruses.

**Figure 2.**
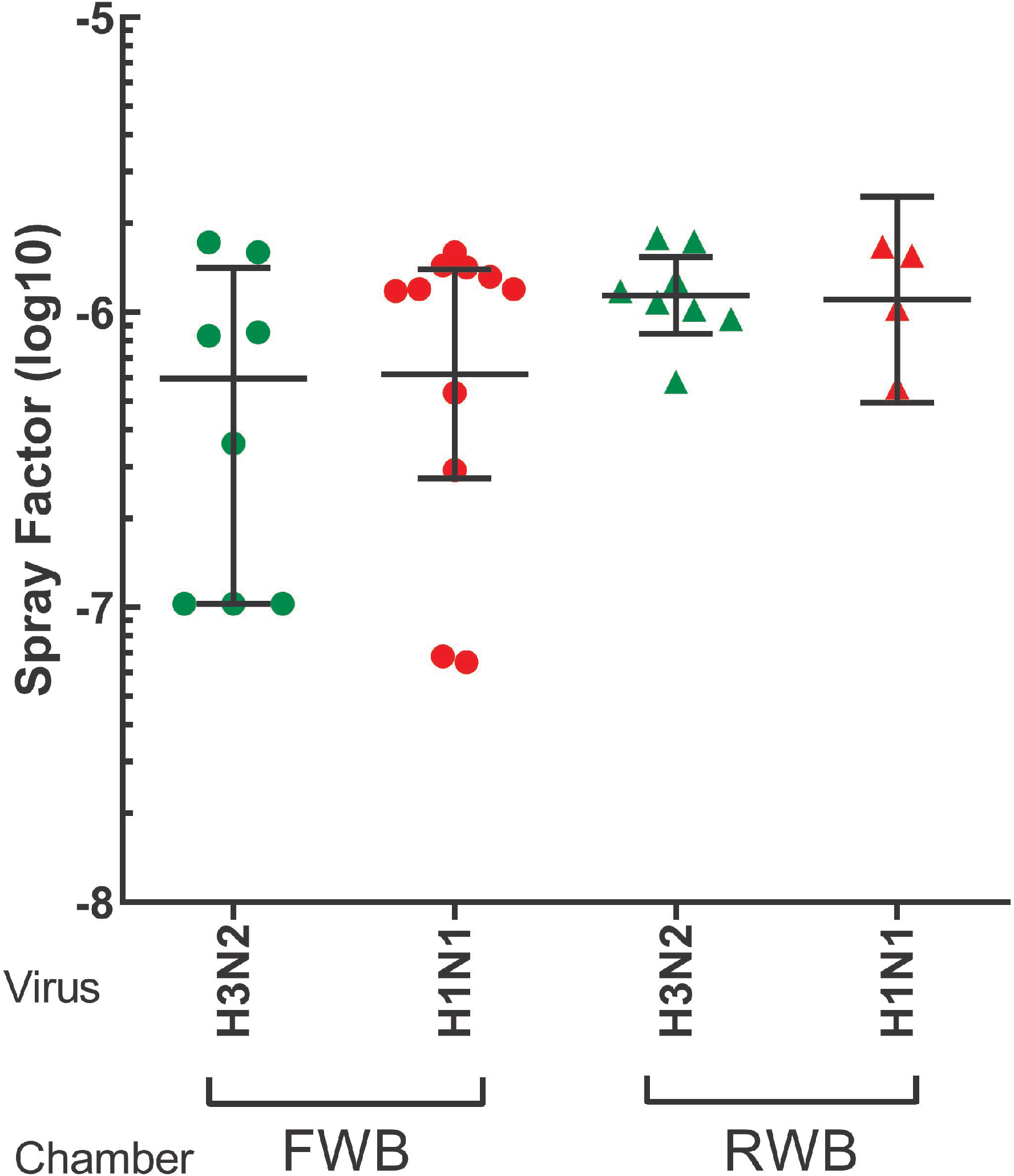
SF does not vary between influenza A strains or exposure chambers. H3N2 and H1N1 influenza viruses were aerosolized using a Collison nebulizer into either a FWB or RWB exposure chamber. Graph shows SF for each combination of virus and exposure chambers. Values shown are individual aerosol runs along with the mean and standard deviation. None of the results were statistically significantly different from the others as determined by a t-test.

**Table 1.** Aerosol performance of influenza strains using the Collison and Aeroneb in different exposure chambers. Shown is the SF of fluorescein salt (the comparator), the SF of influenza, the log reduction in SF between fluorescein and influenza, and aerosol efficiency. Average relative humidity is included to show differences in SF are most likely not due to this factor.

| **Chamber** | **Aerosol Generator** | **Median Fluorescence SF** | **Median Influenza SF** | **Log Reduction** | **Median Influenza AE** | **Average RH (%)** |
| --- | --- | --- | --- | --- | --- | --- |
| **NOT** | **Collison** | **3.44E-06** | **4.34E-06** | **-0.10** | **13.87%** | **85.83** |
|  | **Aeroneb** | **2.68E-06** | **1.18E-06** | **0.36** | **2.10%** | **87.22** |
| **FWB** | **Collison** | **6.35E-06** | **1.20E-06** | **0.72** | **6.84%** | **61.21** |
|  | **Aeroneb** | **6.73E-06** | **5.80E-06** | **0.06** | **31.91%** | **66.64** |
| **NHP HP** | **Collison** | **5.07E-06** | **1.13E-06** | **0.65** | **5.35%** | **81.15** |
|  | **Aeroneb** | **9.97E-06** | **7.52E-06** | **0.12** | **32.27%** | **97.95** |

**Figure 3.**
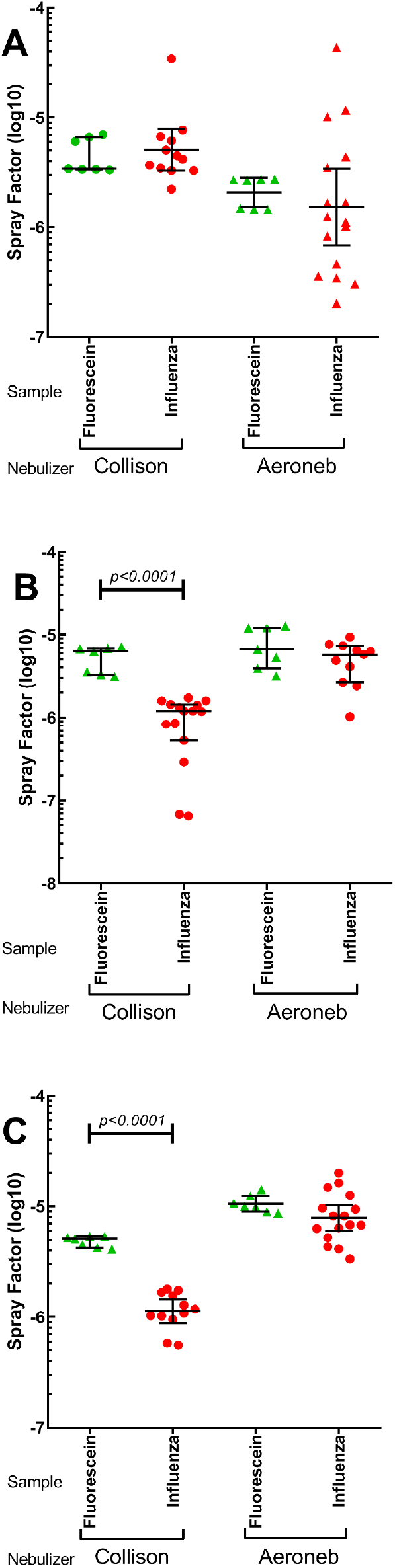
Better aerosol performance using the Aeroneb with influenza viruses. Graphs show the SF of fluorescein salt and influenza in the A) NOT, B) FWB, and C) NHP HO chambers using the Collison or the Aeroneb. Values shown are individual aerosol runs with mean and standard deviation. Black horizontal bars indicate results that are statistically different between fluorescein salt and influenza SF, determined using a t-test with Welch’s correction, with the *p* value shown above the bar.

Comparison of aerosol performance of influenza strains between the Collison and Aeroneb was assessed in the rodent nose-only tower (NOT), the FWB chamber, and the NHP HO chamber. In the NOT, the SF for influenza was higher with the Collison and this difference was significant (*p* = 0.0145) (Table 1, Figure 3A). The range of influenza SF generated by the Aeroneb in the NOT was also substantially broader than was seen with other nebulizer/chamber combinations (coefficient of variation = 2.09). Fluorescein was added as a control to measure impact of the two nebulizers on pathogen viability. For both the Collison and Aeroneb in the NOT, there was little or no loss when comparing influenza SF to fluorescein SF. In contrast, in both the FWB and NHP HO chambers the Aeroneb outperformed the Collison as measured by SF and AE (*p*<0.0001) (Table 1). Further, there was a significant decrease in SF between the fluorescein salt and influenza with the Collison in both the FWB and NHP HO chambers (*p*<0.0001 for both) (Figure 3B-C). This drop was not seen with the Aeroneb, suggesting there is considerable loss of viable influenza in aerosols generated by Collison but not the Aeroneb in these chambers.

To evaluate whether these results were specific to influenza viruses, we also generated aerosols of RVFV into a RWB chamber and compared results obtained with the Aeroneb to prior data obtained with the Collison. As shown in Figure 4A, the Aeroneb did generate a higher SF of RVFV and the improvement was statistically significant (*p*<0.0001). For the encephalitic alphaviruses, the Aeroneb was used to generate aerosols in an NHP HO chamber (Figure 4B). For all three viruses, using the Aeroneb generated a SF that was a ½-1 log_10_ improvement in SF over similar results obtained previously with the Collison *(D.S. Reed, personal observation),* however those results with the Collison were obtained with different virus isolates with some differences in viral plaque assays and media so the results are not directly comparable.

**Figure 4.**
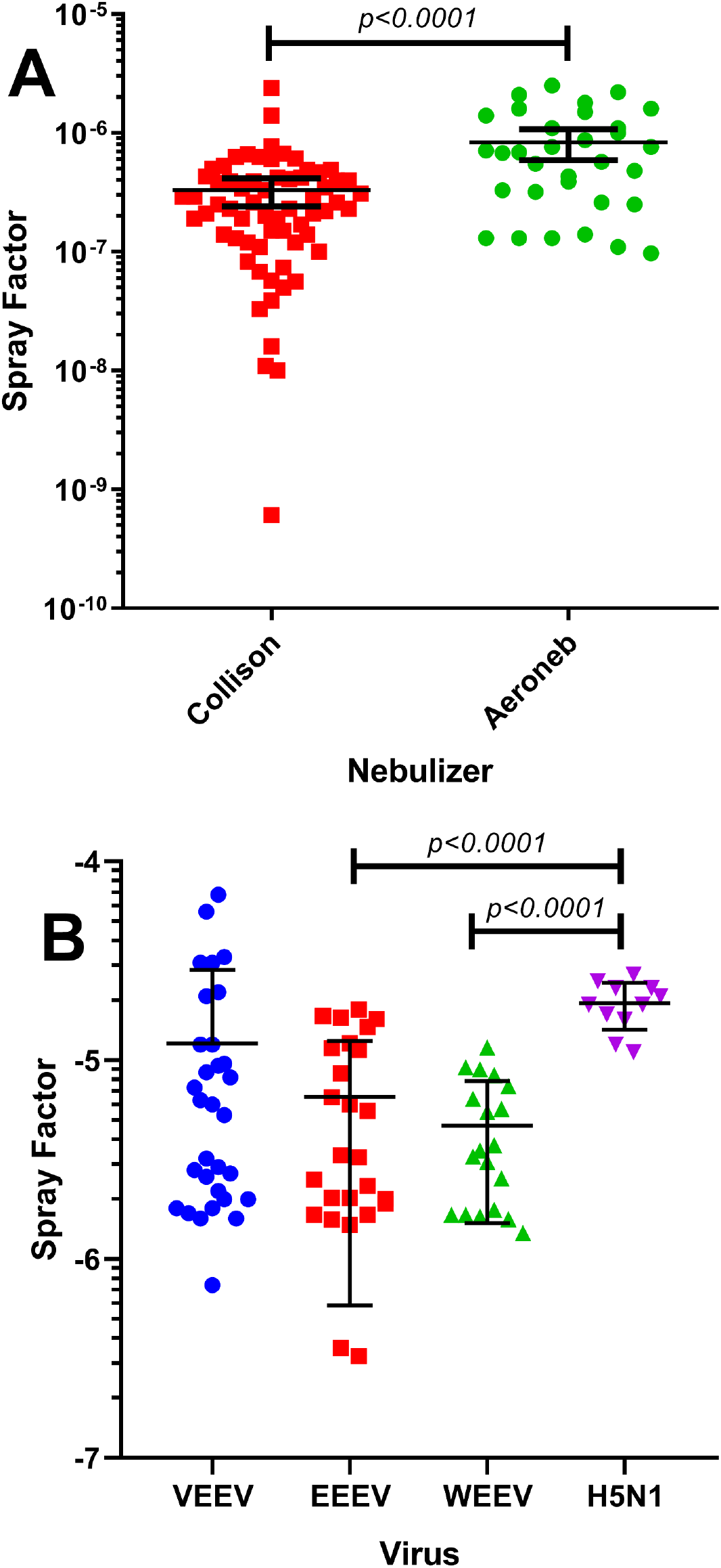
Aerosol performance of the Aeroneb with other viral pathogens. Graphs show the SF of A) RVFV with the Collison and the Aeroneb and B) encephalitic alphaviruses and H5N1 using the Aeroneb. Values shown are individual aerosol runs with mean and standard deviation. Black horizontal bars indicate results that are statistically different between fluorescein salt and influenza SF, determined using a t-test with Welch’s correction, with the p value shown above the bar.

### Aerosolization of *vegetative gram-negative bacteria*

After demonstrating the dramatic improvement in SF for viral pathogens with the Aeroneb, we sought to determine whether similar improvements would be seen with a vegetative bacterium. We had previously shown no difference in SF between attenuated (LVS, the live vaccine strain) and virulent (SCHU S4) strains of *F. tularensis* (Faith et al, 2012). In those studies, we also found that the broth media used to propagate *F. tularensis* greatly impacted SF, as did the relative humidity in the chamber. Aerosol performance of Brain-Heart Infusion (BHI)-grown LVS with the Aeroneb and Collison was assessed in the NOT and the RWB chambers without supplemental humidification. In the NOT, the Collison generated a better SF and higher AE for LVS than did the Aeroneb; this difference was significant (*p*< 0.0001) (Table 2, Figure 5). In contrast, in the RWB chamber, the Aeroneb had a slightly better SF than the Collison which also was significant (*p*=0.0004). AE was also higher for the Aeroneb than the Collison in the RWB. When comparing LVS SF to fluorescein SF in the NOT, we saw a significant, 2 to 3 log_10_ decrease in the SF of LVS with both the Collison and the Aeroneb (*p*=0.0079, 0.0006, respectively) (Figure 5A). An even more substantial decrease in the LVS SF compared to the fluorescein SF was seen in the RWB chamber for both nebulizers (*p*<0.0001 for both) (Figure 5B). This would suggest both nebulizers cause considerable loss of viable LVS although the impact is less in the NOT. This is likely due to the high relative humidity (RH) achieved in the NOT. The higher RH generated by the Aeroneb in the RWB could also explain the superior LVS SF obtained with the Aeroneb in that chamber, as we have previously seen that raising RH above 60% improves LVS SF substantially (3).

**Figure 5.**
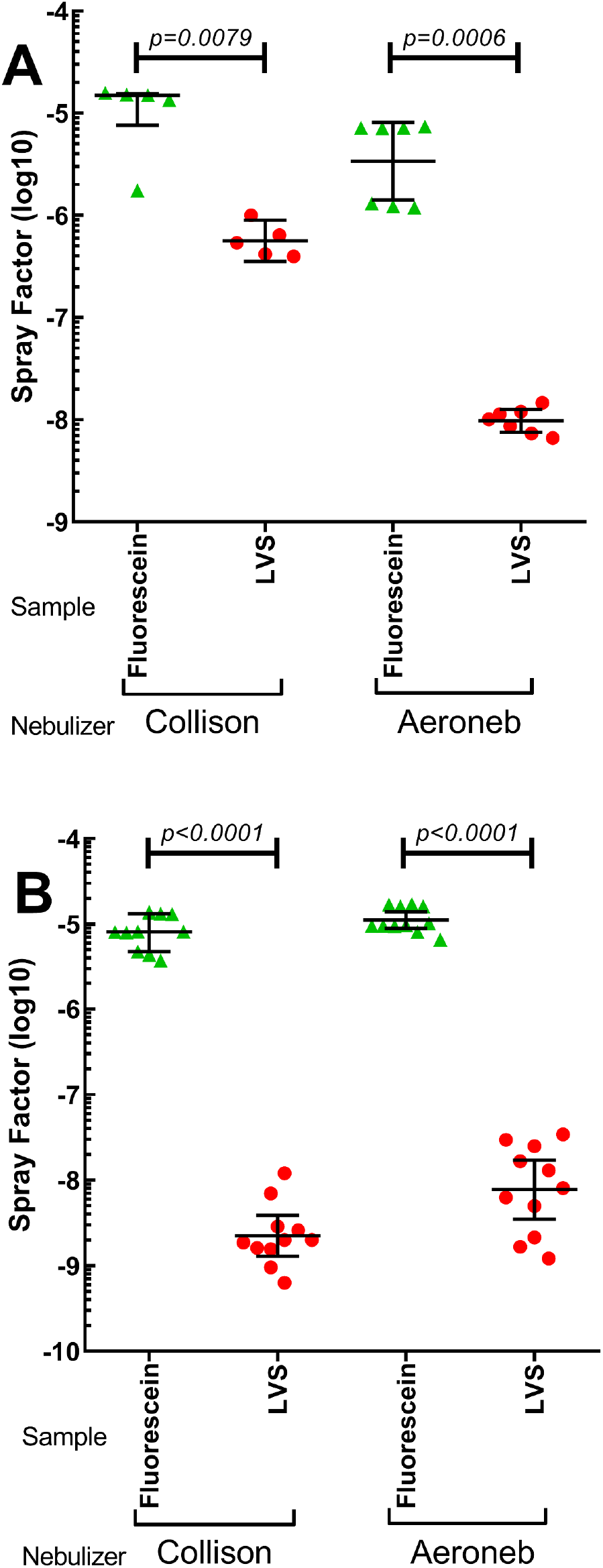
The Collison resulted in better aerosol performance than the Aeroneb. Shown is the SF of fluorescein salt and LVS in the A) NOT and B) RWB using the Collison or the Aeroneb. Values shown are individual aerosol runs with mean and standard deviation. Black horizontal bars indicate results that are statistically different between fluorescein salt and influenza SF, determined using a t-test with Welch’s correction, with the p value shown above the bar.

**Table 2.** Aerosol performance of LVS strains using the Collison and Aeroneb in different exposure chambers. Shown is the SF of fluorescein salt (the comparator), the SF of LVS, the log reduction in SF between fluorescein and LVS, and aerosol efficiency for the different generator and exposure chambers tested. Average relative humidity is included to show differences in SF are most likely not due to this factor.

| **Chamber** | **Aerosol Generator** | **Median Fluorescence SF** | **Median Influenza SF** | **Log Reduction** | **Median Influenza AE** | **Average RH (%)** |
| --- | --- | --- | --- | --- | --- | --- |
| **NOT** | **Collison** | **1.50E-05** | **5.4E-07** | **1.44** | **1.598%** | **82.56** |
|  | **Aeroneb** | **7.08E-06** | **1.01E-08** | **2.85** | **0.035%** | **78.81** |
| **RWB** | **Collison** | **8.14E-06** | **2.01E-09** | **3.61** | **0.012%** | **58.33** |
|  | **Aeroneb** | **9.74E-06** | **8.12E-09** | **3.08** | **0.108%** | **69.21** |

### Particle sizes generated by the Collison and Aeroneb

The Collison has been shown to generate a small (1-2 μm MMAD) particle that is relatively monodisperse. These particles would reach the lower respiratory tract, including the alveolar regions. The information from the manufacturer of the Aeroneb indicates it would generate a somewhat larger particle (average 3.1 μm) which should also reach the alveolar regions. Using an APS 3321, we evaluated particle sizes generated by the Collison and Aeroneb in the different chambers. Initially, we used small (400 or 900 nm) microspheres, however, the Aeroneb was not able to generate good, consistent aerosols with these microspheres. We believe that this difficulty was a result of the microspheres clumping and not being able to readily pass through the vibrating mesh, however, mild sonication did not measurably improve the results (data not shown). If larger particles cannot readily pass through the vibrating mesh, this may contribute to the lower SF obtained with LVS with the Aeroneb. For this reason, we used fluorescein instead of microspheres to measure particle size. The results are shown in Table 3. Particle sizes obtained for the Collison were larger than expected, which we believe may be due to higher surface tension in the aerosolized particles caused by the fluorescein salt. What table 3 does show though is that except for the NOT, the Aeroneb consistently generated larger particles than the Collison and with a broader distribution (as measured by GSD) in all of the chambers tested. The Aeroneb also generated a higher humidity in each chamber tested except for the NOT, which would at least partly explain the differences in particle size seen. Even with the larger particle sizes obtained with the Aeroneb using fluorescein, between 70-80% of the particles measured were ≤5 μm MMAD. The only nebulizer/chamber combination to achieve less than 70% was the Collison in the NOT, which only had 55.97% of particles ≤5 μm. This larger particle size is likely a result of the higher humidification achieved in the NOT by the Collison.

**Table 3.** Particle size generated using the Collison and Aeroneb to aerosolize fluorescein salt in different exposure chambers. Shown is the mass mean aerodynamic diameter (MMAD), geometric standard deviation (GSD), count median aerodynamic diameter (CMAD), and percentage of particles less than or equal to 3.5 and 5 μm in size, and absolute humidity (A.H.) in the exposure chamber, in g/m^3^.

| Chamber | Nebulizer | MMAD | GSD | CMAD | % $\leq 3.5$<br>$\mu\text{m}$ | % $\leq 5$ $\mu\text{m}$ | A.H. |
| --- | --- | --- | --- | --- | --- | --- | --- |
| NHP | Collison | 3.05 | 3.67 | 0.67 | 60.00 | 84.06 | 1018.05 |
|  | Aeroneb | 3.93 | 2.37 | 0.67 | 44.84 | 71.57 | 1256.05 |
| RWB | Collison | 2.96 | 2.84 | 0.65 | 64.07 | 89.11 | 822.88 |
|  | Aeroneb | 3.05 | 4.53 | 0.58 | 60.59 | 84.88 | 1039.48 |
| NOT | Collison | 5.05 | 1.54 | 0.63 | 27.47 | 55.97 | 1360.18 |
|  | Aeroneb | 3.05 | 2.94 | 0.58 | 59.03 | 80.89 | 1302.00 |
| FWB | Collison | 2.21 | 3.41 | 0.63 | 70.34 | 86.55 | 677.36 |
|  | Aeroneb | 3.28 | 2.36 | 0.67 | 54.86 | 74.46 | 921.15 |

## Discussion

Aerosol performance can be affected by a variety of different factors, from pre-aerosolization factors, such as pathogen growth conditions, to post-processing factors, such as concentration determination (3, 15). Thus, prior to beginning aerosol studies with animal models, it is important to characterize and understand the impact of aerosol equipment selection, pathogen handling techniques, and environmental parameters on the reproducibility of a research design. The Collison 3-jet nebulizer has long been used as the “gold standard” for infectious disease aerobiology studies because of its ease of use and relatively monodisperse particle size that can reach the deep lung of rodents, ferrets, rabbits and nonhuman primates. However, the method by which aerosols are generated by the Collison have been considered ‘harsh’ and could damage microorganisms, impacting the dose required to cause infection/disease and the host response to infection (12). The Collison also requires a relatively high volume of challenge material (10 ml), which can be difficult to generate depending on the agent and nebulizer concentration needed to achieve a desired challenge dose. These deficiencies can be a substantial impediment to aerosol studies, particularly for pathogens that require a high challenge dose to achieve infection/disease (e.g., alphaviruses in macaques). Alternative nebulizers that generate small particles that would penetrate to the deep lung (≤5 μm), are less harsh on the microorganism being aerosolized, and require less challenge material to achieve comparable or higher doses would be desirable.

In agreement with what we have reported previously for *F. tularensis* and RVFV, the choice of exposure chamber impacts aerosol performance with smaller chambers (by total volume) typically producing a better SF than a larger chamber. The data we report here also demonstrate that while the choice of nebulizer does affect SF, the impact is dependent upon the chamber used. In the NOT, the Aeroneb did not improve SF compared to the Collison for either LVS or influenza. Yet in the RWB, FWB, and NHP HO chambers, the Aeroneb dramatically improved SF performance compared to the Collison for influenza and other viral pathogens but had minimal impact on the SF for LVS. Particle sizes generated by the Aeroneb were generally larger than those generated by the Collison, except in the NOT, but 70-80% of the particles generated by the Aeroneb were in the ‘respirable’ range (≤ 5 μm MMAD) that would reach the deep lungs. Humidity levels were generally higher with the Aeroneb compared to the Collison, except in the NOT, which may explain the differences seen in SF and particle sizes with the Aeroneb in the other chambers.

Another important difference to note between the Collison and the Aeroneb is the volume needed for aerosolization and total volume aerosolized. The Collison requires 10 ml of sample for aerosol generation, while the Aeroneb requires 5-6 ml for a 10-minute aerosol. On average, the Collison aerosolized 3ml of sample while the Aeroneb aerosolized 4ml of sample during that 10-minute aerosol. Additional challenge material could be added to the Aeroneb for aerosol exposures longer than 10 minutes; while technically feasible for the Collison, this would not be easily done. For each generator/exposure chamber setup, aerosol efficiency correlated with the SF, indicating that the improvement in SF for the Aeroneb compared to the Collison in the RWB, FWB and NHP HO chambers was not due to the increased volume of material aerosolized by the Aeroneb.

Prior studies have suggested the Collison may damage pathogens during the process of aerosolization through mechanical and shear forces (12). Fluorescein salt was used in some experiments to act as a surrogate for microorganisms to determine the ideal SF of each aerosol generator given loss within the system. The small size and lack of a membrane ensures the fluorescein salt will not be damaged by the aerosolization process, and thus the loss of fluorescein salt during aerosolization can be attributed to leaks in the exposure system and adhesion of aerosol particles to equipment. Any additional decrease in SF of pathogens compared to the SF for fluorescein following aerosolization is likely due to loss of viability in the organism. The vegetative LVS bacteria had a significant drop in SF relative to fluorescein salt (1-3 log_10_) in all the combinations of nebulizer and exposure chamber tested here. This was despite the relatively high RH generated by either nebulizer. Reflecting the apparent loss in bacterial viability, the AE was quite low for LVS using either nebulizer.

Influenza SF also dropped relative to fluorescein salt for the Collison in the FWB and NHP HO chamber but not in the NOT. Surprisingly, the SF for influenza aerosolized with the Aeroneb did not drop relative to fluorescein salt in any of the chambers tested. This data would suggest that for viral pathogens, the superior SF of the Aeroneb to the Collison may be at least partially due to improved aerosol viability. Relative humidity did not appear to substantially alter SF for influenza although the RH was high in all the nebulizer/chamber combinations tested. Additional data generated with RVFV and the encephalitic alphaviruses further confirmed the superior SF performance of the Aeroneb with viral pathogens. The data presented in this paper indicate the Aeroneb is a suitable alternative to the Collison for infectious disease aerobiology research, particularly for viral pathogens. This data has been successfully used in developing a macaque model for respiratory exposure to highly pathogenic avian influenza (16). Exploration of aerosol generators other than the Collison is recommended when evaluating new animal models for human respiratory infections.

## Materials and Methods

### Animal Use

Experiments described in this report that involved animals were approved by the University of Pittsburgh’s IACUC. Research was conducted in compliance with the Animal Welfare Act Regulations and other Federal statutes relating to animals and experiments involving animals and adheres to the principles set forth in the Guide for Care and Use of Laboratory Animals, National Research Council, 1996. The University of Pittsburgh is accredited by the Association for the Assessment and Accreditation of Laboratory Animal Care (AAALAC).

### Biosafety

All aerosol experiments for this study were performed in a class III biological safety cabinet within the dedicated Aerobiology Suite inside a Biosafety Level-3 (BSL-3) facility operated by the Center for Vaccine Research. For respiratory protection during H5N1, alphavirus, or Rift Valley Fever virus experiments, personnel wore powered air purifying respirators (PAPRs) while performing plaque assays within class II biosafety cabinets at BSL-3 conditions, using Vesphene IIse (diluted 1:128, Steris Corporation) for disinfection. Spatial and temporal separation was maintained between H5N1, Rift Valley Fever, and all other infectious agents. Work with *F. tularensis* LVS strain and seasonal influenza was conducted at BSL2+ conditions in a class II biosafety cabinet using 10% bleach or Vesphene IIse (1:128) for disinfection.

### Bacteria

A frozen stock of Live Vaccine Strain (LVS) *F. tularensis* originally obtained from Jerry Nau and passaged a single time in culture were used for aerosol experiments. Prior to aerosol exposure, LVS was grown on Cysteine Heart Agar (CHA; BD Difco™ and BD BBL™, Becton Dickinson, La Jolla, CA) for two days at 37°C, 5% CO_2_ and then overnight in Brain Heart Infusion (BHI) broth (BD BBL^TM^) supplemented with 2.5% ferric pyrophosphate and 1.0% L-Cysteine hydrochloride as previously described (3). Cultures were incubated at 37°C in an orbital shaker at 200rpm and harvested between 15 to 18 hours to ensure the bacteria were in the logarithmic growth phase. The O.D. of the culture was read and bacterial concentration estimated based on previously determined OD to CFU ratios (3). The concentration of LVS was confirmed by colony counts on CHA.

### Influenza

Two H1N1 strains, A/PR/8/34 obtained from Rich Webby, and A/Ca/04/09 from the Biodefense and Emerging Infectious Resources were used in these experiments. An H3N2 virus (influenza A/Syd/5/37) obtained from Michael Murphy Corb and an H5N1 virus (A/Vietnam/1203/04) obtained from Daniel Perez were also used. The H1N1 and H3N2 viruses were propagated in MDCK cells and frozen at −80 until use. The H5N1 (A/Vietnam/1203/2004) stock was propagated in SPF chicken eggs and the stock frozen at −80 until use. Temporal and spatial separation of all strains of influenza was maintained throughout the experiments; H1N1 and H3N2 viruses were used at BSL-2 while H5N1 was used at BSL-3. Prior to aerosol experiments, influenza viral stocks were diluted in viral growth media (Dulbecco’s Modified Eagle’s Medium, 2.5% of 7.5% bovine serum albumin fraction V, 1% penicillin/streptomycin, 1% HEPES buffer, and 0.1% TPCK trypsin).

### Rift Valley Fever virus (RVFV)

The stock of RVFV (isolate ZH501) used in these experiments was derived from an infectious clone as previously described (17). Prior to aerosol experiments, it was thawed and diluted in DMEM containing 2% FBS, glycerol and Antifoam A for aerosolization as previously described.

### Alphaviruses

Venezuelan equine encephalitis virus (VEEV; isolate INH9813), western equine encephalitis virus (WEEV; isolate Fleming), and eastern equine encephalitis virus (EEEV; isolate V105) were derived from infectious clones of human isolates passaged a single time in BHK cells. Stocks were thawed and diluted in Optimem for aerosolization.

### TCID50

confluent MDCK cells (ATCC CCL-34) were infected with tenfold serial dilutions of influenza samples in a 96-well plate. The plates were incubated at 37°C/5% CO_2_ for 48 hours. Cells were then examined under a microscope for cytopathic effect (CPE) as compared to the uninfected MDCK cell controls. Each well was scored as positive or negative for CPE. Viral titers were then calculated using the method described by Reed and Muench.

### Plaque assay

virus samples were adsorbed onto confluent monolayers of Vero, Vero E6, or MDCK cells in duplicate wells of a 6-well plate for one hour at 37°C/5% CO_2_. After incubation, inoculum was removed and cells were overlaid with a 1% nutrient overlay (2X Modified Eagle Medium, BSA, penicillin/streptomycin, 2% agarose). Plates were incubated at 37°C/5% CO_2_ for up to 5 days, depending on virus. Cells were fixed with 37% formaldehyde, agar plugs were removed, and cells were stained with a 0.1% crystal violet stain to visualize plaques. Wells with 15 to 100 plaques were counted for titer calculations. H5N1 plaque assays were performed in the same manner as seasonal influenza plaque assays with the following changes: following the addition of inoculum, the plates were incubated at 4°C for 10 minutes, then incubated at 37°C/5% CO2 for 50 minutes; a 0.9% nutrient overlay was used instead of a 1.0% nutrient overlay.

### Aerosol Exposures

The AeroMP or Aero 3G aerosol management systems (Biaera Technologies, Hagerstown, MD) were used to control, monitor, and record aerosol parameters during aerosol experiments. Unless otherwise noted, aerosols were ten minutes in length. The airflow parameters of the aerosol experiments were programmed based on chamber volume in accordance with protocols used to infect animals. Air input (primary and secondary air) and vacuum (exhaust and sampler) were set in balance at one-half of the chamber volume, to insure one complete air change in the exposure chamber every two minutes. Aerosols were generated using either a 3-jet Collison nebulizer or an Aeroneb nebulizer (see Figure 1). Airflow through the Collison was set at 7.5 lpm and 26-30 psi. The Aeroneb utilizes a vibrating mesh, not pressurized air, for generating aerosol particles. The Aeroneb was placed in line with the secondary/dilution air to push the air into the exposure chamber. Because exposure chamber structure and volume can influence aerosol performance (5), four exposure chambers were used for these experiments: the rodent nose-only tower (NOT), the rodent whole-body chamber (RWB), the ferret whole-body chamber (FWB), and the nonhuman primate head only chamber (NHP HO), with chamber volumes of 12L, 39L, 44L, and 32L respectively.

### Aerosol Sampling

Bioaerosol sampling was performed using the all glass impinger (AGI; Ace Glass, Vineland, NJ) calibrated with the Gilibrator to ensure an airflow of 6.0 ± 0.25 L/min. The AGI is attached to the side of the aerosol exposure chamber in an area close to the breathing zone. For LVS aerosols, 10ml of BHI broth and 40μl of antifoam A (Fluka, cat. #10794) was added to each AGI. For virus aerosols, 10ml of cell culture media and 80μl of antifoam was added to each AGI. For RVFV aerosols, glycerol was also added. For VEEV, WEEV, and EEEV aerosols, 1% FCS was also added to the culture media. Aerosol concentration was determined as previously described (3, 5).

### Aerosol Performance

Aerosol performance between nebulizers was compared using SF and aerosol efficiency (AE). SF was determined as previously described (3, 5), the ratio of the aerosol concentration (determined from the AGI) to the starting concentration in the aerosol generator. AE is the ratio of the aerosol concentration to the theoretical maximum aerosol concentration as previously described (18). Aerosol particle size as measured by mass median aerodynamic diameter (MMAD) and geometric square deviation (GSD) using an aerodynamic particle sizer (APS) model #3321 (TSI, Shoreview, MN).

### Fluorescein

Fluorescein salt (Sigma) was added to some aerosol experiments to be used as an indicator of maximum SF given natural loss. Fluorescein salt was dissolved at a concentration of 0.1mg in 1ml of ddH2O prior to addition to nebulizer contents. Initial studies were conducted (data not shown) to verify that addition of fluorescein did not alter pathogen viability or quantitation in culture, whether by plating on agar (*F. tularensis*) or TCID50/plaque assay (influenza).

### Statistical analysis

GraphPad Prism^®^ 6 was used to create all figures and to perform two-sided Mann-Whitney U tests to compare the SF and aerosol efficiency between nebulizers. This nonparametric test was chosen due to the non-normal distribution of results and the high frequency of outliers.

## Acknowledgements

The research described herein was sponsored by the National Institute of Allergy & Infectious Diseases at the National Institutes of Health, grants R01 A102966-01A1 & R21 NS088326-01 and the Defense Threat Reduction Agency Grant #W911QY-15-1-0019 and is sponsored by the Department of the Army, U.S. Army Contracting Command, Aberdeen Proving Ground, Natick Contracting Division, Ft. Detrick Maryland. Any opinions, findings, and conclusions or recommendations expressed in this material are those of the author(s) and do not necessarily reflect the position or the policy of the Government and no official endorsement should be inferred. Special thanks to past and present members of the Hartman and Klimstra labs at the University of Pittsburgh, especially Stacey Barrick, Matthew Dunn, Dr. Christina Gardner, Theron Gilliland, Jr., Joe Albe, and Aaron Walters.

## References

1. Fernstrom A, Goldblatt M. 2013. Aerobiology and its role in the transmission of infectious diseases. Journal of pathogens 2013.

2. Thomas R, Davies C, Nunez A, Hibbs S, Flick-Smith H, Eastaugh L, Smither S, Gates A, Oyston P, Atkins T. 2010. Influence of particle size on the pathology and efficacy of vaccination in a murine model of inhalational anthrax. Journal of medical microbiology 59:1415–1427.

3. Faith S, Smith LK, Swatland A, Reed DS. 2012. Growth conditions and environmental factors impact aerosolization but not virulence of Francisella tularensis infection in mice. Frontiers in cellular and infection microbiology 2:126.

4. Saini D, Hopkins GW, Chen C-j, Seay SA, Click EM, Lee S, Hartings JM, Frothingham R. 2011. Sampling port for real-time analysis of bioaerosol in whole body exposure system for animal aerosol model development. Journal of pharmacological and toxicological methods 63:143–149.

5. Reed DS, Bethel LM, Powell DS, Caroline AL, Hartman AL. 2014. Differences in aerosolization of Rift Valley fever virus resulting from choice of inhalation exposure chamber: implications for animal challenge studies. Pathogens and disease 71:227–233.

6. Cox C, Goldberg L. 1972. Aerosol survival of Pasteurella tularensis and the influence of relative humidity. Applied microbiology 23:1–3.

7. Hood A. 1977. Virulence factors of Francisella tularensis. Epidemiology and Infection 79:47–60.

8. Roy C, Reed D, Hutt J. 2010. Aerobiology and inhalation exposure to biological select agents and toxins. Veterinary pathology 47:779–789.

9. Lackemeyer MG, Kok-Mercado Fd, Wada J, Bollinger L, Kindrachuk J, Wahl-Jensen V, Kuhn JH, Jahrling PB. 2014. ABSL-4 aerobiology biosafety and technology at the NIH/NIAID integrated research facility at Fort Detrick. Viruses 6:137–150.

10. May K. 1973. The Collison nebulizer: description, performance and application. Journal of Aerosol Science 4:235–243.

11. Roy C, Pitt L. 2012. Infectious disease aerobiology, Biodefense Research Methodology and Animal Models, 2nd ed. Taylor & Francis Group.

12. Zhen H, Han T, Fennell DE, Mainelis G. 2014. A systematic comparison of four bioaerosol generators: Affect on culturability and cell membrane integrity when aerosolizing Escherichia coli bacteria. Journal of Aerosol Science 70:67–79.

13. Thomas RJ, Webber D, Hopkins R, Frost A, Laws T, Jayasekera PN, Atkins T. 2011. The cell membrane as a major site of damage during aerosolization of Escherichia coli. Applied and environmental microbiology 77:920–925.

14. Zhang G, David A, Wiedmann TS. 2007. Performance of the vibrating membrane aerosol generation device: aeroneb micropump nebulizer™. Journal of Aerosol Medicine 20:408–416.

15. Goodlow RJ, Leonard FA. 1961. Viability and infectivity of microorganisms in experimental airborne infection. Bacteriological reviews 25:182.

16. Wonderlich ER, Swan ZD, Bissel SJ, Hartman AL, Carney JP, O’Malley KJ, Obadan AO, Santos J, Walker R, Sturgeon TJ. 2017. Widespread virus replication in alveoli drives acute respiratory distress syndrome in aerosolized H5N1 influenza infection of macaques. The Journal of Immunology:1601770.

17. Bales JM, Powell DS, Bethel LM, Reed DS, Hartman ALJFic, microbiology i. 2012. Choice of inbred rat strain impacts lethality and disease course after respiratory infection with Rift Valley Fever Virus. 2:105.

18. Dabisch P, Yeager J, Kline J, Klinedinst K, Welsch A, Pitt ML. 2012. Comparison of the efficiency of sampling devices for aerosolized Burkholderia pseudomallei. Inhalation toxicology 24:247–254.

